# Activation of a *Vibrio cholerae* CBASS anti-phage system by quorum sensing and folate depletion

**DOI:** 10.1101/2023.04.04.535582

**Authors:** Geoffrey B. Severin, Miriam S. Ramliden, Kathryne C. Ford, Andrew J. Van Alst, Ram Sanath-Kumar, Kaitlin A. Decker, Brian Y. Hsueh, Soo Hun Yoon, Lucas M. Demey, Brendan J. O’Hara, Christopher R. Rhoades, Victor J. DiRita, Wai-Leung Ng, Christopher M. Waters

**Author notes:** Corresponding Author: 5180 Biomedical and Physical Sciences 567 Wilson Road, East Lansing, MI 48824. Current Address, Department of Microbiology and Immunology, University of Michigan, Ann Arbor, Michigan, USA. Declarations of interest: none.

## Abstract

A major challenge faced by bacteria is infection by bacteriophage (phage). Abortive infection is one strategy for combating phage in which an infected cell kills itself to limit phage replication, thus protecting neighboring kin. One class of abortive infection systems is the <u>c</u>yclic oligonucleotide <u>b</u>ased <u>a</u>nti-phage <u>s</u>ignaling <u>s</u>ystem (CBASS) which relies on two core enzymatic activities; an oligo-nucleotide cyclase that is activated following phage infection and a cyclic-oligo-nucleotide sensitive effector whose activity kills the infected cell. However, the mechanisms behind the deployment and activation of these lethal CBASS systems prior-to and following infection have largely remained a mystery. While exploring unique genomic features of the current pandemic *Vibrio cholerae* biotype El Tor for clues underlying its pandemic success we found its CBASS was spuriously activated by the folate biosynthesis inhibitor sulfamethoxazole, but only after the population had reached a high-cell density. This population density dependent activity revealed that transcription of both the oligo-nucleotide cyclase, *dncV*, and the CBASS phospholipase effector, *capV*, is enhanced at high-cell density by quorum sensing. Together, these results demonstrate that the *V. cholerae* CBASS is deployed when the environment is densely populated and activated in response to a perturbation in folate biosynthesis.

**Significance:** To counteract infection with phage, bacteria have evolved a myriad of molecular defense systems. Some of these systems initiate a process called abortive infection, in which the infected cell kills itself to prevent phage propagation. However, such systems must be inhibited in the absence of phage infection to prevent spurious death of the host. Here we show that the <u>c</u>yclic oligonucleotide <u>b</u>ased <u>a</u>nti-phage <u>s</u>ignaling <u>s</u>ystem (CBASS) accomplishes this by sensing intracellular folate molecules and only expressing this system in a group. These results enhance our understanding of the evolution of the 7^th^ *V. cholerae* pandemic and more broadly how bacteria defend themselves against phage infection.

## Introduction

The diarrheal disease cholera, caused by the Gram-negative bacterium *Vibrio cholerae*, is spread through consumption of contaminated food and water (1). Of the seven recorded cholera pandemics the classical biotype is believed to have caused at a minimum both the 5^th^ (1881 - 1896) and 6^th^ (1899 - 1923) pandemics whereas the El Tor biotype is responsible for initiating and perpetuating the 7^th^ pandemic (1961 - today) (2, 3). While strains of the classical biotype are now rarely encountered in environmental and clinical settings, numerous assays have been developed to help distinguish *V. cholerae* isolates as belonging to either the classical or El Tor biotypes (4, 5). These include disparate growth on citrate, hemolytic activity, casein proteolysis, production of acidic or neutral byproducts from growth on glucose, differences in virulence gene allele sequences and expression, and disparate sensitivities to the cationic antimicrobial peptide polymyxin B and the folate biosynthesis inhibitor sulfamethoxazole (SMX).

Two of the largest genetic differences between the *V. cholerae* biotypes are the Vibrio Seventh Pandemic Islands 1 and 2 (VSP-1 and 2) genomic islands that are defining features of the El Tor biotype and absent in the classical biotype (3, 6, 7). VSP-1 and 2, which collectively contain ∼36 genes, are hypothesized to have played a critical role in the pandemic evolution of the El Tor biotype and many recent studies have begun to explore the biological functions they encode. The predominant function of these islands appears to be protection against invasive biological elements as two phage defense systems, AvcID (8, 9) and a Type II <u>c</u>yclic oligonucleotide <u>b</u>ased <u>a</u>nti-phage <u>s</u>ignaling <u>s</u>ystem (CBASS) (10–13), are encoded on VSP-1 while VSP-2 encodes the *ddmABC* operon, which inhibits plasmid acquisition, plasmid stability, and phage invasion (14, 15). In addition to biological defense, a three gene operon on VSP-2 that is encoded in some strains of El Tor mediates aerotaxis in response to zinc (16). Outside of these four systems, little is known of the function of the VSP islands or their contribution to the emergence of the El Tor biotype.

The VSP-1 CBASS encompasses a four gene operon [*vc0178(capV)-vc0179(dncV)*-*vc0180(cap2)-vc0181(cap3)*]. In this system, via an unknown mechanism, phage infection stimulates DncV, a member of the CD-NTase family of enzymes (17), to synthesize of 3’3’ cyclic GMP-AMP (cGAMP) (11, 18). cGAMP then allosterically activates the phospholipase CapV, which rapidly degrades the infected cell’s own membrane (10). Cap2 enhances the production of cGAMP by post translationally modifying the C-terminus of DncV in a manner analogous to ubiquitination (12, 13). Conversely, Cap3 suppresses DncV activity by precisely proteolyzing this same C-terminal modification (12, 13). Ultimately, activation of VSP-1 CBASS rapidly kills the infected cell to restrict phage propagation and protect neighboring kin from further phage predation, a mechanism called abortive infection (11). While CBASS systems are widely encoded in bacterial genomes (11, 17, 19), we are just beginning to learn how these lethal population-level phage defense systems are controlled.

Bacteria sense their population density through quorum sensing (QS). Based on the constituency and abundance of bacteria in the environment, a bacterium uses QS to enact gene expression patterns that promote or discourage participation in coordinated population-level behaviors (reviewed in (20, 21)). Bacteria assess the local population density and composition by producing, secreting, and detecting small molecules called auto-inducers (AIs) whose environmental concentrations are a proxy for bacterial abundance. In *V. cholerae*, QS gene expression programs for low-cell density (LCD) and high-cell density (HCD) are regulated by the transcription factors LuxO and HapR, respectively. At LCD, when the environmental concentrations of AIs are low, LuxO is active and *hapR* mRNA is degraded. As AI concentrations increase in the environment, a proxy for an increasing population density, LuxO activity is inhibited via dephosphorylation, driving *hapR* expression and enabling induction of the HCD gene expression regulon. The dichotomous QS-dependent activities of LuxO and HapR are primarily responsible for the differential regulation of more than 500 *V. cholerae* genes (22). These population-dependent changes in transcription determine whether *V. cholerae* pursue behavioral strategies best suited for a solitary or group lifestyle at LCD or HCD.

To better understand the contribution of the VSP islands to the evolution of the 7^th^ *V. cholerae* pandemic, we explored if the VSP islands in the El Tor biotype were responsible for some of the well-known phenotypic differences between strains of the classical and El Tor biotypes. We report that El Tor’s sensitivity to the folate biosynthesis inhibitor SMX is dependent on the VSP-1 encoded Type-II CBASS. This sensitivity results from the spurious activation of DncV and the subsequent activation of the phospholipase CapV. Furthermore, during these studies we found that the expression of *dncV* and *capV* are induced at HCD by the *V. cholerae* QS pathway, consistent with its function as a population-level phage defense mechanism. Our findings identify both transcriptional and post-transcriptional mechanisms that lead to the deployment and activation of the El Tor *V. cholerae* CBASS.

## Results

### VSP-1 and -2 do not impact metabolic phenotypes commonly used to distinguish the El Tor and classical *V. cholerae* biotypes

We examined whether the VSP-1 and VSP-2 islands contributed to phenotypic behaviors commonly attributed to the El Tor biotype by performing several biotyping assays comparing classical strain O395 (23) and the El Tor strain C6706str2 (C6706) (24) with those of single VSP island mutants (ΔVSP-1 and ΔVSP-2) and a double VSP island mutant (ΔVSP1/2). O395 is known to be deficient in both protease and hemolysin production in comparison to C6706, as demonstrated on casein milk agar and blood agar plates, respectively (4). Simple streaks of these strains on milk agar and sheep blood agar confirmed differential degradation of casein and hemolytic activities between O395 and C6706, as indicated by the absence and presence of a zone of clearing on each medium, respectively (Figs. 1A & 1B). We found that deletion of one or both VSP islands did not affect either activity as all three mutants phenocopied the parental C6706 strain (Figs. 1A & 1B). Similarly, classical O395 cannot utilize citrate as a sole carbon source (25) or grow on MacConkey agar, while C6706 can do both. Again, we found that the VSP islands do not contribute to these two phenotypes as all three mutants grew comparably to the parental C6706 strain on these media (Figs. 1C and 1D). Finally, El Tor strains are known to produce acetoin upon fermentation of glucose while classical strains do not (26). Using the colorimetric Voges-Proskauer assay to measure acetoin by the generation of a red color, we found supernatants of C6706 produced a deep red color, indicative of acetoin production, while classical O395 was weakly pink (Fig. 1E). Production of acetoin was not grossly impacted upon deletion of either or both VSP islands (Fig. 1E) as these strains also produced a deep red color like the parental C6706. While casein degradation in milk agar and production of the fermentative bioproduct acetoin have been linked to QS (4, 27) and the inability of O395 to perform these functions is likely due, in-part, to its non-functional *hapR* allele (28), we nonetheless demonstrate the VSP islands do not contribute to these phenotypes in C6706.

**Figure 1:**
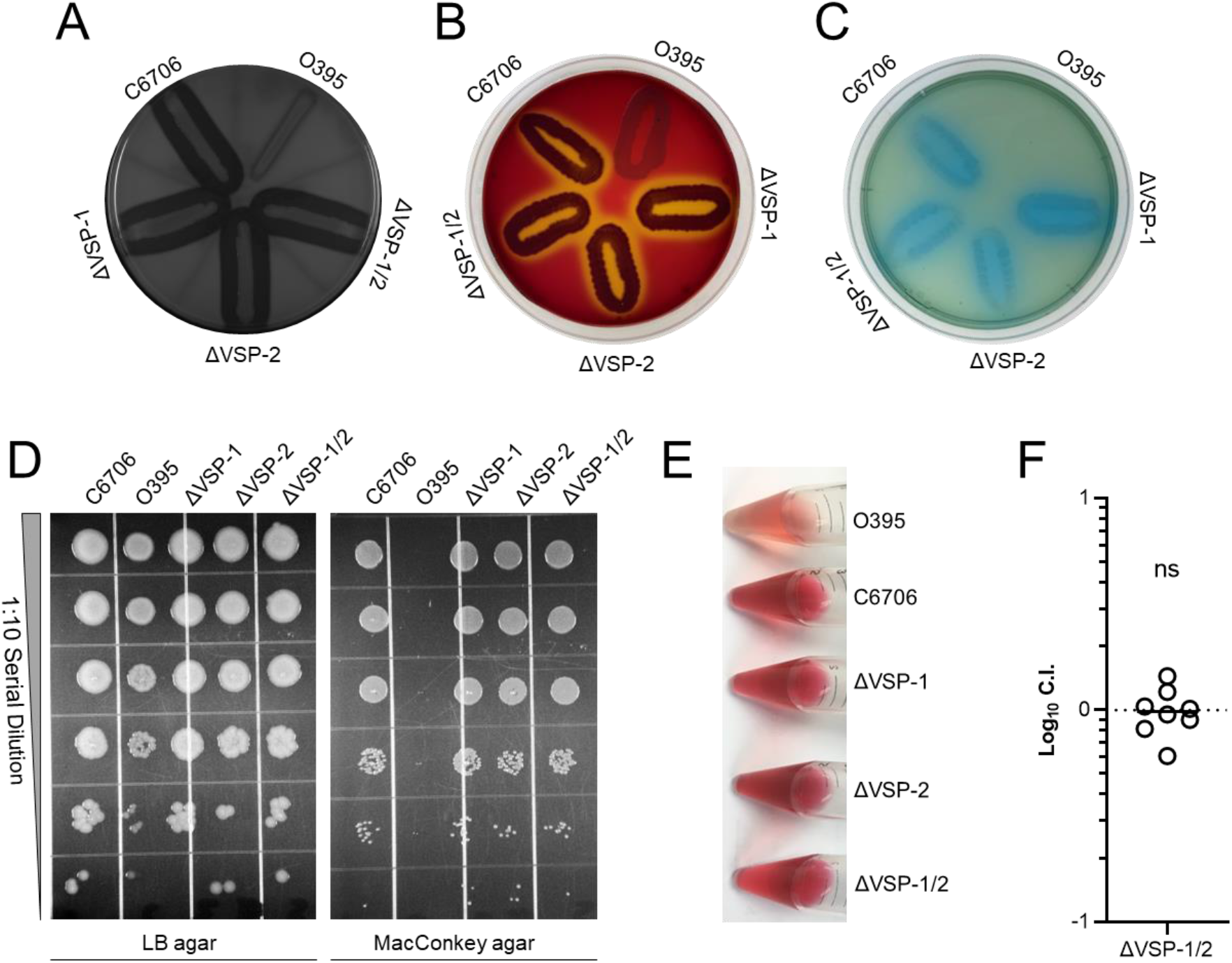
The VSP islands do not contribute to common *V. cholerae* biotyping phenotypes or virulence in a murine model of cholera. Differential biotyping phenotypes between classical O395, El Tor C6706, and C6706 VSP island mutants (ΔVSP-1, ΔVSP-2, and ΔVSP-1/2) demonstrating strain specific (**A**) proteolysis of casein on milk agar, (**B**) hemolytic activity on blood agar, (**C**) growth on citrate minimal medium agar, (**D**) growth on matched LB and MacConkey agar plates, and (**E**) production of acetoin detected by Voges-Proskauer Assay. All images are representative of three independent experimental replicates. (**F**) In vivo competition between a 1:1 mixture of C6706 ΔVSP1/2 and C6706 Δ*lacZ* in an infant mouse model of cholera. Intestinal colony forming units (CFU) were enumerated using blue-white screening 20 hours after oral gavage. N = 8 mice and statistical significance was determined using a one sample *t* test and a hypothetical mean Log_10_ C.I. = 0. The hypothetical mean is represented by a dotted line. The calculated mean Log_10_ C.I. is represented by a solid line. ns = not significant.

### The VSP-1 and -2 islands do not contribute to El Tor C6706 colonization in an infant mouse model

Subtle differences in the sequences of the virulence gene alleles *ctxB* and *tcpA* between strains of the El Tor and classical biotypes have been used for biotyping novel *V. cholerae* isolates (4). The classical biotype is also more permissive in its in vitro expression of the *V. cholerae* virulence regulon compared to the El Tor biotype (29), which anecdotally causes less severe cholera in humans (30). To determine whether the VSP islands collectively contribute to in vivo colonization we performed a competition infection between C6706 and ΔVSP-1/2 in the infant mouse model of cholera. Despite the previous attribution of *dncV*, the CBASS cGAMP synthase located in VSP-1, to colonization (18), we found no competitive defect in the ability of the double VSP island mutant to colonize the infant mouse intestinal tract (Fig. 1F). The discrepancy between this finding and *Davies et al.* 2012 may be attributed to unknown epistatic relationships between *dncV* and other VSP encoded genes which obscure this colonization defect in our study, genetic differences between laboratory lineages of C6706 (31), or specific conditions not replicated in our study. Nevertheless, our result suggests that the collective impact of the VSP islands on colonization in the infant mouse model of cholera is negligible.

### VSP-1 increases sensitivity to sulfamethoxazole

In addition to different metabolic and virulence characteristics, different susceptibilities to antibiotics have also been observed between classical and El Tor strains. One such antibiotic is the cationic antimicrobial peptide polymyxin B, which disrupts the outer membrane of Gram-negative bacteria. Indeed, as previously observed in pandemic *V. cholerae* biotypes (32, 33), we found that classical O395 was more sensitive to polymyxin B than El Tor C6706 with half maximal inhibitory concentrations (IC_50_) of 0.6 and 18.4 µg/mL, respectively (Fig. S1, Table S1). However, deletion of VSP-1, VSP-2, or both islands did not grossly impact El Tor’s susceptibility to polymyxin B (Fig. S1, Table S1) demonstrating the VSP islands do not contribute to this biotype specific phenotype.

In contrast to polymyxin B, strains of the El Tor biotype have been shown to exhibit greater sensitivity to sulfamethoxazole (SMX) than those of the classical biotype (5). SMX impairs folate biosynthesis by inhibiting the activity of dihydropteroate synthase. After treating our strains with a concentration gradient of SMX and measuring culture optical densities after 24 hours we found C6706 was profoundly more sensitive to SMX (IC_50_ 36.5 µg/mL) than classical O395 (IC_50_ 230.7 µg/mL) (Figs. 2A & S2A, Table S1). Surprisingly, both ΔVSP-1 and ΔVSP-1/2 phenocopied O395’s SMX resistance while the ΔVSP-2 mutant retained the parental C6706 SMX sensitivity (Figs. 2A & S2A, Table S1). Hypothesizing SMX sensitivity could be attributed to VSP-1, we reintroduced VSP-1 into the ΔVSP-1/2 mutant on a single copy cosmid (pVSP-1) and found this was sufficient to restore SMX sensitivity (Figs. 2B & S2B). Additionally, provision of pVSP-1 to both O395 and *Escherichia coli* BL21(DE3) also increased each strain’s sensitivity to SMX (Figs. 2B, S2C & S2D). Together, these results demonstrate the disparity in *V. cholerae* biotype specific sensitivities to SMX is the result of a factor encoded on VSP-1.

**Figure 2:**
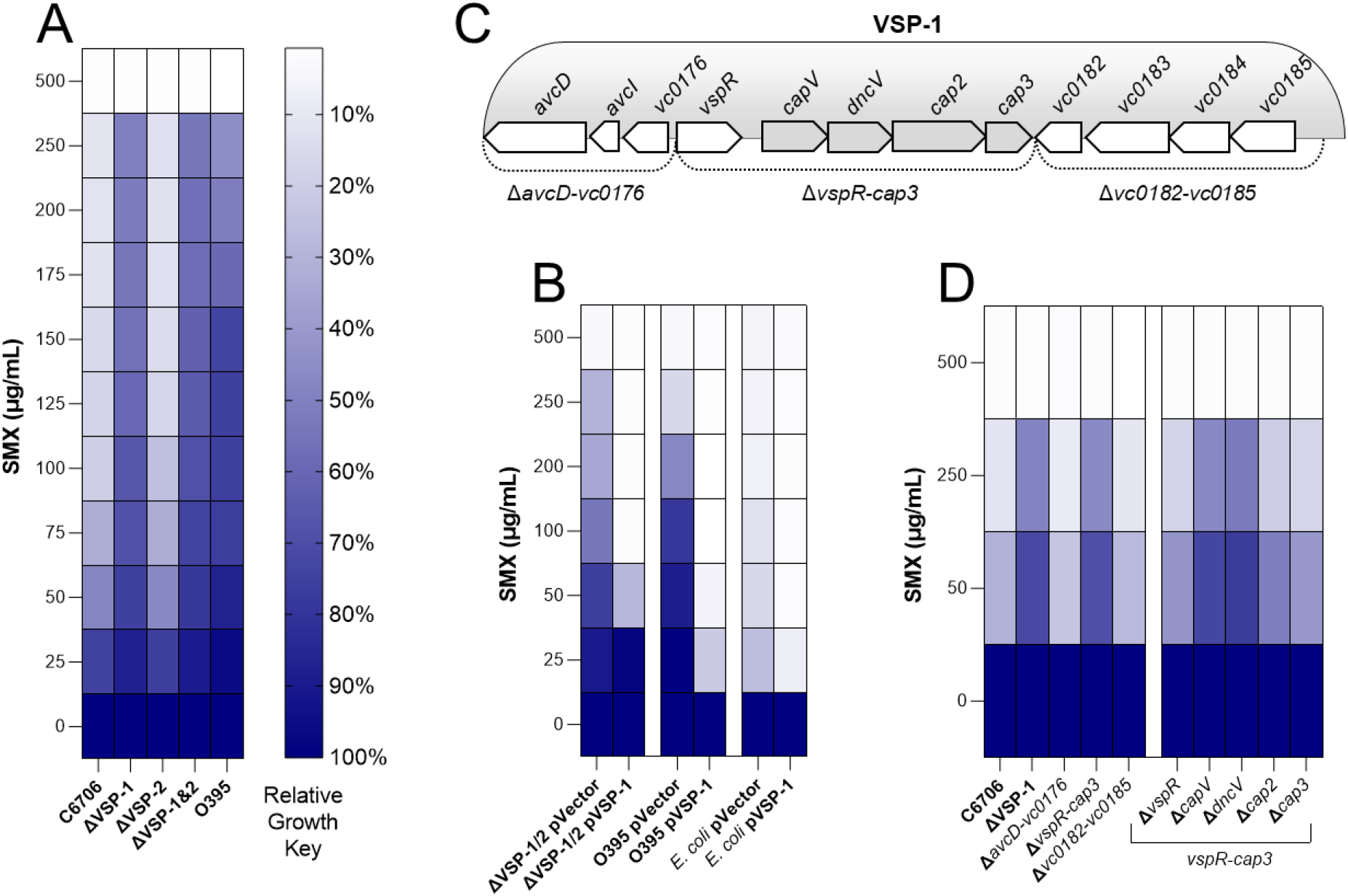
VSP-1 encoded CBASS is responsible for *V. cholerae* biotype specific SMX sensitivity. 24-hour planktonic antibiotic sensitivity assays were performed in a variety of SMX concentration gradients. Heatmaps (**A**), (**B**), and (**D**) represent relative growth for each strain calculated using culture optical densities and the equation (OD_600_ SMX treatment / OD_600_ untreated) and reported as a color-coded mean % of N = 3 biological replicates. The IC_50_ for all strains in (**A**) are presented in Supplementary Table 1. Scatter plots corresponding to (**A**) (**B**) and (**D**) are presented in (Figs. S2A-F). (**C**) Cartoon depiction of the VSP-1 genomic island. Dotted lines indicate partial VSP-1 truncations. Grey chevrons highlight the four gene VSP-1 CBASS operon.

### Sulfamethoxazole sensitivity requires the CBASS genes *dncV* and *capV*

The VSP-1 island of *V. cholerae* C6706 contains 12 known or predicted genes (Fig. 2C). To identify which genes are responsible for SMX sensitivity, we challenged three partial VSP-1 island mutants (Δ*avcD-vc0176*, Δ*vspR-cap3*, and Δ*vc0182-vc0185*) with a gradient of SMX concentrations and found only the Δ*vspR-cap3* mutant was more resistant to SMX (Figs. 2D & S2E). The five genes missing in this mutant include the negative transcriptional regulator of CBASS, *vspR* (18), and the four gene CBASS (*dncV*-*cap3*) (11) (Fig. 2C). It was previously shown that folates are allosteric regulators of DncV which suppress cGAMP synthesis in vitro (34) and that cGAMP is required for the activation of the lethal phospholipase CapV (10). We therefore hypothesized that inhibition of folate biosynthesis by SMX could alleviate the folate-dependent repression of DncV activity and lead to CapV-dependent cell death. In support of this hypothesis, deletion of either *dncV* or *capV* was sufficient to enhance resistance to SMX in these conditions while loss of *vspR*, *cap2*, or *cap3* was not (Figs. 2D & S2F).

### Sulfamethoxazole activates DncV cGAMP synthesis

It was previously reported that the addition of sulfonamide to *E. coli* strains ectopically expressing *dncV* from a high-copy plasmid enhanced the catalytic activity of DncV (34). To test whether SMX treatment might induce cGAMP synthesis by DncV in its native cellular and genetic contexts, we back-diluted stationary phase cultures of C6706 and Δ*capV* 1:1,000 in fresh medium, allowed them to recover for approximately one hour, introduced 100 µg/mL SMX or a DMSO control, and measured culture optical density over the course of six hours. Concurrent with monitoring culture density, we also measured the intracellular concentration of cGAMP using UPLC-MS/MS, but this measurement was only performed in the Δ*capV* cultures where detectable levels of cGAMP can accumulate without inducing CapV-dependent cell lysis (10). Surprisingly, C6706 cultures treated with SMX only exhibited a growth defect after two hours of exposure to the antibiotic (Fig. 3A). This growth defect is a consequence of CapV activity as there was no difference in growth between Δ*capV* cultures treated with and without SMX (Fig. 3A). Similarly, cGAMP was only found in Δ*capV* cultures challenged with SMX and was not detected until more than two hours after its addition (Fig. 3A). While these results demonstrate that SMX stimulates DncV synthesis of cGAMP in its native environment, they also reveal a profound delay exists between introduction of the antibiotic and evidence of DncV activity under these experimental conditions.

**Figure 3:**
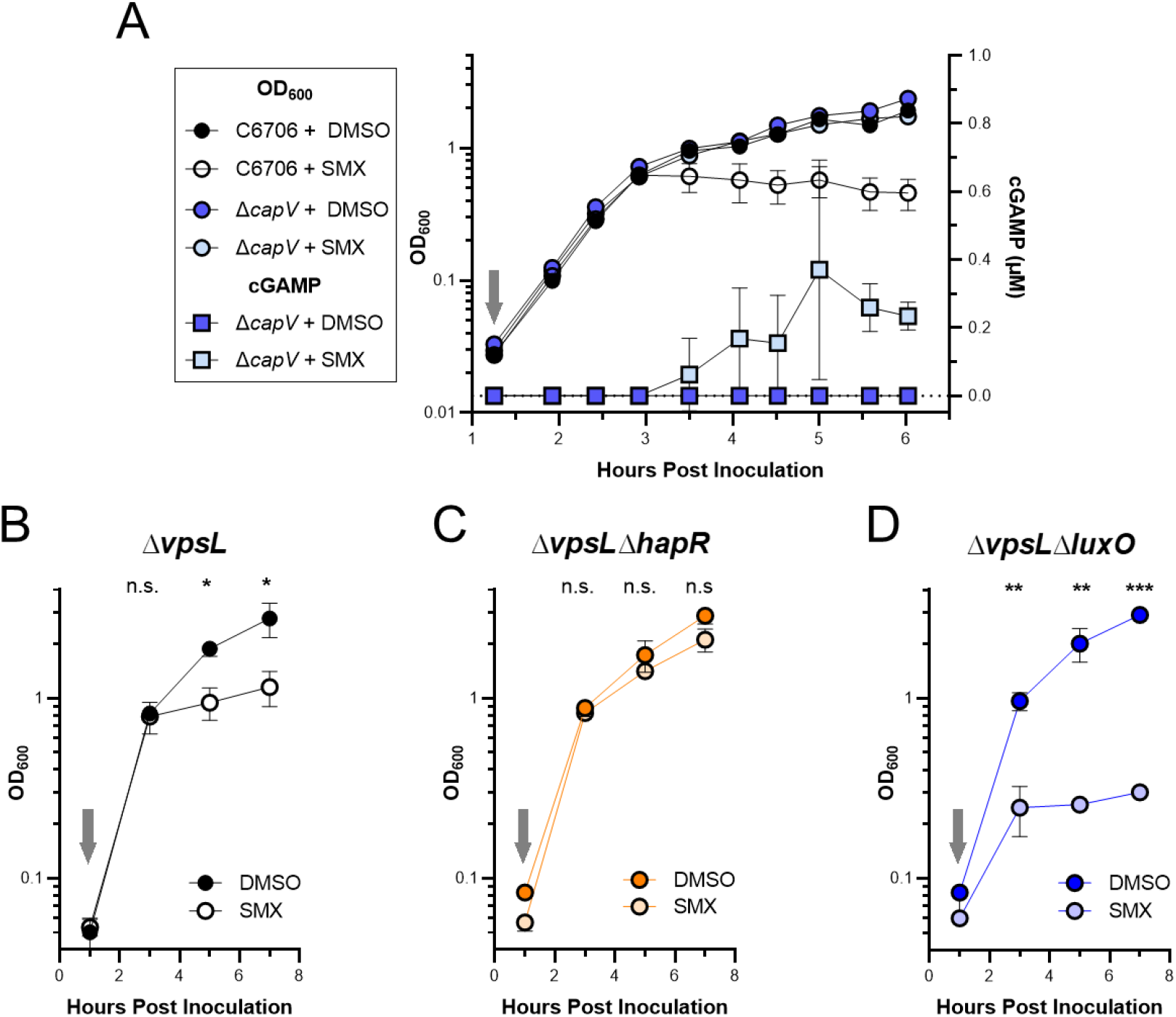
Culture density-dependent sensitivity to SMX reveals quorum sensing regulation of VSP-1 CBASS. (**A**) Growth curves (OD_600_, left y-axis) of WT C6706 and Δ*capV* cultures treated without (+ DMSO) and with 100 µg/mL SMX (+ SMX). Intracellular cGAMP (µM, right y-axis) measured by UPLC-MS/MS in the SMX treated and untreated Δ*capV* cultures. Growth curves of (**B**) quorum fluent Δ*vpsL*, (**C**) LCD-locked Δ*vpsL*Δ*hapR*, and (**D**) HCD-locked Δ*vpsL*Δ*luxO* cultures treated without (+ DMSO) and with 100 µg/mL SMX (+ SMX). Grey arrows indicate addition of 100 µg/mL SMX or DMSO. N = 3 biological replicates and error bars represent standard deviation. Statistical significance calculated using an unpaired *t* test with the Holm-Šídák method (*P < 0.05, **P < 0.005, ***P <0.0005), n.s. = not significant.

### QS contributes to CBASS-dependent SMX sensitivity

Hypothesizing the two-hour delay in SMX-dependent growth inhibition and cGAMP synthesis (Fig. 3A) was an indication of CBASS regulation by QS, we challenged QS mutants locked in LCD or HCD gene expression with 100 µg/mL SMX or a DMSO control and monitored culture densities over time. Because LCD QS mutants have a propensity to form biofilms, which could interfere with measuring optical density, we utilized QS mutants derived from the biofilm-null C6706 Δ*vpsL* background (35). Importantly, there is no difference between C6706 and Δ*vpsL* when challenged with SMX or DMSO in these conditions (Fig. S3). As previously seen for C6706 (Fig. 3A), the Δ*vpsL* cultures with an intact QS system experienced a similar two-hour delay in growth inhibition in response to SMX (Fig. 3B). In agreement with our hypothesis that QS induces CBASS, the LCD-locked Δ*vpsL*Δ*hapR* strain was tolerant of SMX (Figs. 3C) while the HCD-locked Δ*vpsL*Δ*luxO* was hypersensitive to SMX with an abbreviated temporal delay in growth inhibition (Fig. 3D). Together, these results support the argument that QS contributes to the regulation of CBASS.

### QS regulates transcription of the *V. cholerae* CBASS operon

To determine if QS regulation of CBASS was at the level of transcription induction, we first monitored the abundance of *capV* and *dncV* transcripts in cultures of *V. cholerae* C6706 over time using RT-qPCR. Three hours following inoculation and corresponding with ∼ 0.6 OD_600_, a culture density consistent with the transition from LCD to HCD (36), the abundance of each transcript abruptly increased ∼8-fold relative to their initial levels and remained high for the duration of the experiment (Fig. 4A). Notably, the increased abundance of both *capV* and *dncV* transcripts three hours post inoculation precedes both the SMX-dependent detection of cGAMP in the Δ*capV* mutant and growth inhibition by SMXof quorum fluent strains in previous experiments (Figs. 3A, 3B, S3A).

**Figure 4:**
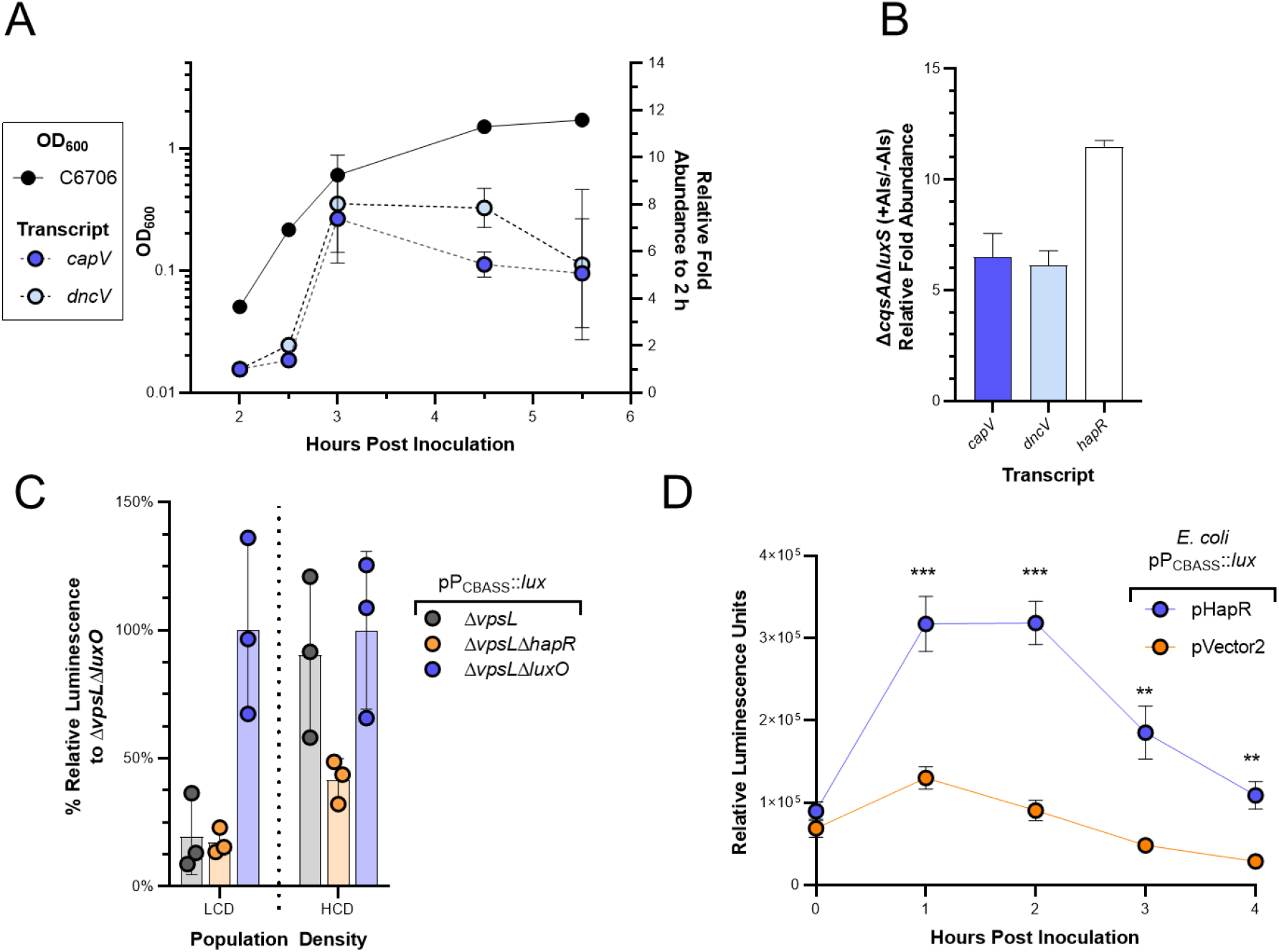
QS regulates expression of *V. cholerae* CBASS. (**A**) Growth curve (OD_600_, left y-axis, solid line) of *V. cholerae* C6706 and the corresponding fold-change in transcript abundance (right-axis) of *capV* and *dncV*, relative to the initial 2-hour time-point, measured using RT-qPCR. N = 2 biological replicates and error bars represent standard deviation. (**B**) Relative transcript abundance of *capV*, *dncV*, and *hapR* measured by qRT-PCR in Δ*csqA*Δ*luxS* grown in the presence (+) and absence (-) of exogenous autoinducers (AIs). N = 3 biological replicates and error bars represent standard error of the mean. (**C**) % relative luminescence units of the indicated strains normalized to the mean lum/OD_600_ of Δ*vpsL*Δ*luxO* maintaining the luminescent transcriptional reporter pP_CBASS_::*lux.* Population densities are low-cell density (LCD) and high-cell density (HCD). N = 3 biological replicates and error bars represent standard deviation. (**D**) Relative luminescence (lum/OD_600_) of *E. coli* maintaining the luminescent transcriptional reporter pP_CBASS_::*lux* and the P_tac_ inducible *hapR* plasmid (pHapR) or a vector control (pVector2) grown in the presence of 6.25 µM IPTG. N = 3 biological replicates and error bars represent standard deviation. Statistical significance calculated using an unpaired *t* test with the Holm-Šídák method (**P < 0.005, ***P <0.0005), n.s. = not significant.

To confirm the observed increase in *capV* and *dncV* transcripts were the result of HCD gene expression we measured their abundance in a strain of C6706 (Δ*csqA*Δ*luxS*) (37) which is incapable of producing the two primary *V. cholerae* AIs, CAI-2 and AI-1. In monoculture, Δ*csqA*Δ*luxS* is locked in LCD gene expression, regardless of the population density, but can be converted to HCD gene expression by the introduction of exogenous AIs. Using RT-qPCR, we found when Δ*csqA*Δ*luxS* was grown in the presence of exogenous AIs there was at least a 6-fold greater abundance of *capV* and *dncV* transcripts than when grown in their absence (Fig. 4B). As a control, the abundance of *hapR* transcript also increased upon AI addition indicating the cultures had been converted to HCD gene expression (Fig. 4B).

To better understand how QS was regulating the abundance of CBASS transcripts, we constructed a luminescent transcriptional plasmid reporter containing a 913 nt region 5’ of the *capV* translational start site with the CBASS promoter (pP_CBASS_::*lux*) and measured luminescence generated by biofilm-null Δ*vpsL* QS mutants at LCD and HCD. The HCD-locked strain, Δ*vpsL*Δ*luxO* was the most luminescent at both densities whereas the LCD-locked strain, Δ*vpsL*Δ*hapR*, was sparingly luminescent regardless of the culture density (Fig. 4C). The quorum fluent strain, Δ*vpsL*, resembled Δ*vpsL*Δ*hapR* at LCD and Δ*vpsL*Δ*luxO* at HCD (Fig. 4C). These results indicated that transcription initiation of the CBASS promoter is positively regulated at HCD.

The transcription factor primarily responsible for HCD gene expression in *V. cholerae* is HapR (28). To determine if HapR could enhance expression of the CBASS promoter independent of additional *V. cholerae* genes, we measured the luminescent output of pP_CBASS_::*lux* in a heterologous *E. coli* host and provided *hapR* in trans on an inducible plasmid (pHapR). The relative luminescence of *E. coli* cultures maintaining pHapR were significantly greater than the vector control following introduction of the inducer (Fig. 4D). In total, our data indicate CBASS is expressed as part of the HCD regulon in *V. cholerae* C6706 and this expression is induced by HapR.

### HapR-dependent induction of CBASS transcription contributes to phage defense

QS induction of the CBASS operon suggests that this system would exhibit higher levels of phage defense at HCD. However, we have not identified a condition in which the *V. cholerae* CBASS operon protects against infection by the three major *V. cholerae* lytic phage ICP-1, ICP-2, and ICP-3 (38). Thus, we are unable to test this prediction in *V. cholerae* expressing CBASS from its native genomic context. To circumvent this challenge, we examined if the *V. cholerae* CBASS operon provided phage defense when expressed from a low-copy cosmid (pVSP-1) in *E. coli* to the lytic phage T2 in shaking liquid cultures. In these conditions, pVSP-1 alone provided only minimal enhancement of *E. coli* growth when challenged with T2 at low multiplicities of infection (MOIs) (Fig. 5A, S4). However, HapR expression substantially enhanced protection by pVSP-1 to T2 phage infection, demonstrating that QS enhancement of CBASS expression leads to greater phage defense (Fig. 5A, S4).

**Figure 5:**
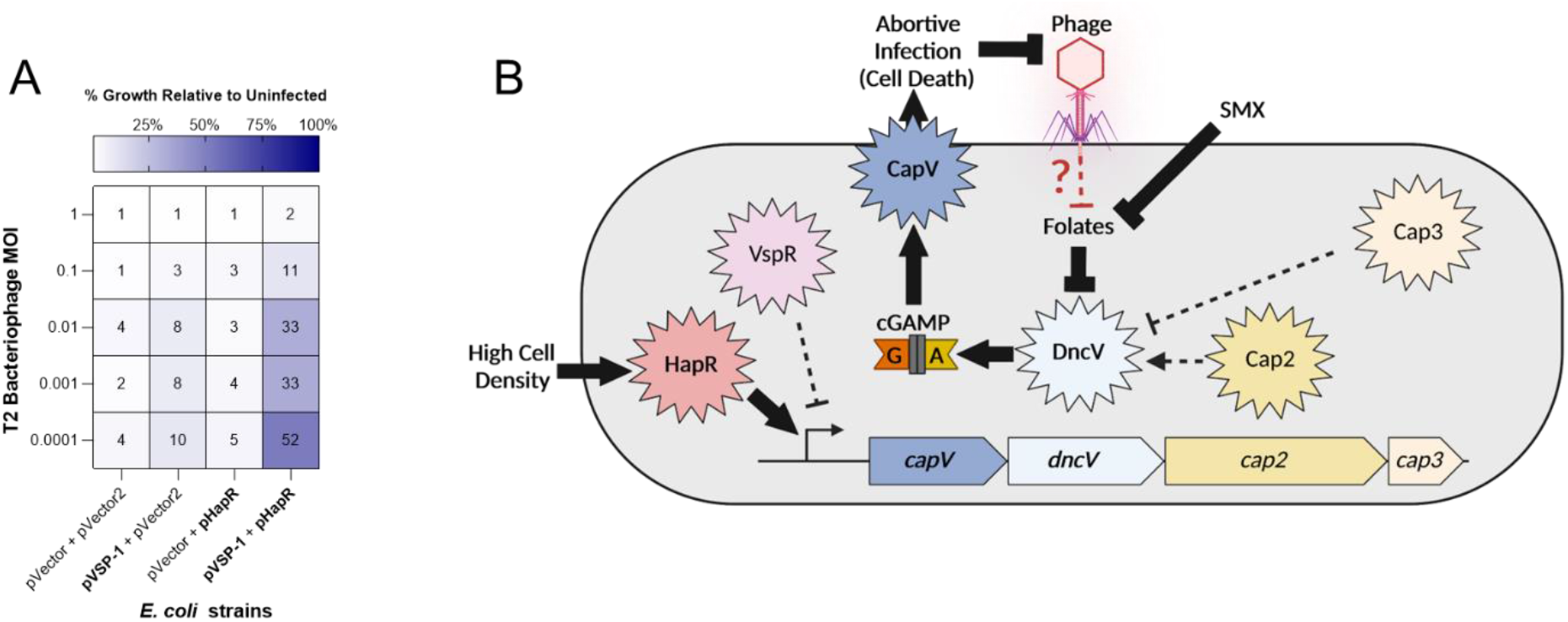
HapR enhances VSP-1 mediated phage defense in *E. coli*. (**A**) Growth of *E. coli* containing either pHapR induced with 10 μM IPTG and pVSP-1 with their associated vector controls after overnight growth with T2 phage is shown as a mean percent of the uninfected culture from N = 3 biological replicates. Mean % presented numerically and by heatmap for each condition. Scatter plot of data presented in (Fig. S4). (**B**) Model of folate (by SMX) and QS (by HapR) regulation of CBASS activity and expression. At HCD, HapR induces transcription of the CBASS operon. Inhibition of folate biosynthesis by SMX alleviates the folate-dependent non-competitive inhibition of DncV leading to synthesis of cGAMP and activation of the phospholipase CapV. CBASS activity ultimately culminates in abortive infection that thwarts phage predation by limiting phage replication. Solid arrows and brakes indicate regulatory mechanisms that influence CBASS activity addressed in this study. Black hatched arrows and brakes indicate mechanisms known to contribute to CBASS activity but were not found to significantly contribute to QS or folate mechanisms illuminated in this study. Red brake represents a hypothesized phage-dependent disturbance in folates which is sensed by DncV to initiate abortive infection by the *V. cholerae* CBASS. Created with BioRender.com.

## Discussion

The earliest strains of the El Tor biotype were first encountered during the period of 1897 – 1938 and considered non-pathogenic enteric commensals (3). Through genotypic analysis of early and modern El Tor strains, Hu *et al.* (3) traced the evolutionary linage of the El Tor biotype from non-pathogenic commensal to pandemic scourge in six phases. While these phases include the predictable acquisition of key virulence factors including *tcpA*, *toxT*, and cholera toxin, the 5^th^ stage (1925-1954) is primarily defined by El Tor’s acquisition of VSP-1 and -2 from unknown origins. While it has been hypothesized that acquisition of VSP-1 and -2 potentiated El Tor’s pandemicity and rise to global dominance in the 7^th^ Pandemic, we are just beginning to understand their utility and the functions they encode.

Our analyses show that VSP-1 and -2 do not contribute to previously identified metabolic differences between El Tor and Classical biotypes nor do they collectively influence colonization in an infant mouse infection model. Rather, we demonstrate that the increased sensitivity to the antibiotic SMX in the model El Tor strain C6706 is due to the spurious activation of the abortive infection anti-phage Type-II CBASS encoded on VSP-1. Upon exposure to SMX, inhibition of folate biosynthesis de-represses DncV which leads to the synthesis of cGAMP (Fig. 5B). This increase of intracellular cGAMP allosterically activates the phospholipase CapV leading to the degradation of bacterial membranes and cell death (10) (Fig. 5B). We note that a recent study independently reached the same conclusion finding that the CBASS system of *V. cholerae* strain N16961 drove sensitivity to SMX (39). N16961 is a natural locked LCD mutant of *V. cholerae* with a non-functional *hapR,* and thus the role of QS in CBASS activity could not be observed in this study. In response to increased usage of SMX to treat cholera infections, SMX resistance has been observed in 79% of O1/O139 isolates (40). However, such resistance is not due to the loss of CBASS function but rather the acquisition of other resistance mechanisms like the SXT integrative conjugative element (41, 42). The fact that El Tor strains retain CBASS function and VSP-1 while acquiring resistance to SMX via alternate mechanisms argues that the VSP-1 island continues to play an important role in the fitness of circulating El Tor biotype strains. One such role could be enhanced resistance to phage predation which has been hypothesized to contributed to the global rise of the El Tor biotype and displacement of the classical biotype prior to the 7^th^ pandemic (43).

The link between SMX exposure and the activation of the VSP-1 CBASS is likely the result of non-competitive inhibition of DncV by folate molecules. The x-ray crystal structure of DncV unexpectedly revealed a molecule of 5-methyl-tetrahydrate folate bound on the opposite face of the active site (34). Further study revealed that the in vitro cyclase activity of purified DncV was inhibited by the addition of various folate molecules (34). Additionally, introduction of the folate biosynthesis inhibitors sulfonamide and trimethoprim to *E. coli* expressing recombinant DncV led to an increase in the intracellular concentration of cGAMP relative to untreated cells (34). DncV belongs to the CD-NTase family of cyclic oligo-nucleotide synthases, which are often allosterically regulated (reviewed in (44)). For example, metazoan cGAS only synthesizes 2’3’ cyclic GMP-AMP when bound to dsDNA as this is a biological signal for viral invasion or genome instability (45, 46) while the homolog cGLR1 from *Drosophila melanogaster* responds to dsRNA (47). Activation of DncV in *V. cholerae* by SMX indicates folates likely allosterically inhibit this CD-NTase in its native host.

Given that DncV initiates an abortive infection program (48), and its activity is inhibited by folates (34), we hypothesize that disruption of folate metabolism is a cellular indication of phage infection. In support of this idea, phage often encode nucleotide biosynthetic enzymes and extensively remodel cellular nucleotide pools, which can in turn alter folate metabolism. For example, the bacteriophage T4 encoded protein 55.1 forms a complex with *E. coli* FolD, a bifunctional enzyme that catalyzes interconversion of 5,10-methylene-tetrahydrofolate and 10-formyl-tetrahydrofolate, and induces hypersensitivity to trimethoprim (49). Additionally, the T4 strain, T4D, has been shown to shift the metabolism of folate compounds during infection in *E. coli* (reviewed in (50)). T4 phage have also been shown in numerous studies to be susceptible to CBASS anti-phage activities (11–13). Our results support a model (Fig. 5B) that depletion of folates upon phage infection is the activation signal for DncV to initiate abortive infection, though this remains to be formally tested. It is also unclear whether folates facilitate or antagonize the dichotomous activities of Cap2 and Cap3 which enhance and suppress DncV activity, respectively, through post-translation modification of its C-terminus (12, 13). Under the conditions we tested, activation by folate depletion is indifferent to the post-translational modification state of DncV as loss of either *cap2* or *cap3* did not substantially affect SMX sensitivity (Figs. 2D & S2F).

Our studies of SMX activation of DncV showed a consistent delay in cell killing, which we determined is due to the QS-dependent transcription of CBASS by the HCD transcriptional regulator HapR (Fig. 5B). In agreement with this, RNA-seq analysis of *V. cholerae* C6706 demonstrated that CBASS transcripts were significantly more abundant at HCD than LCD (22). QS regulation appears to be another mechanism to restrict the activity of the VSP-1 CBASS system to the most appropriate conditions, thereby preventing spurious activation of DncV and CapV and unnecessary cell death. Identification of the VSP-1 encoded transcription factor VspR (Fig. 2C, 5B) was pivotal to the discovery of DncV (18). ChIP analysis revealed VspR was associated with DNA sequences found within the CBASS locus and *vspR* mutants contained a greater abundance of CBASS gene transcripts, implicating it as a negative transcriptional regulator of CBASS (18). Contrary to our expectation, under our experimental conditions *vspR* did not contribute to CBASS-dependent SMX sensitivity (Fig. 2D & S2F). If and how VspR and QS regulation of CBASS overlap remains to be explored.

The role of QS in defense against phage has been previously described (reviewed in (51)) and some notable examples include the regulation of phage receptor abundance in *E. coli* (52) as well as haemaglutinin protease production in El Tor *V. cholerae* (53), CRISPR-Cas expression in *Pseudomonas aeruginosa* (54), and the *hapR* independent HCD-regulation of the El Tor VSP-II encoded *ddmABC* anti-phage and plasmid defense system (15). From an ecological perspective, reserving the expression and activity of the VSP-1 CBASS to situations where El Tor *V. cholerae* would find itself in an environment densely populated with kin fits with its biological function of preventing phage infection by abortive replication. HCD populations are environments where phage will be the most prevalent and bacteria at greater risk of infection. At LCD, when neighbors are scarce, abortive infection is unlikely to be an evolutionarily advantageous strategy and infected cells may instead rely on non-lethal phage defense mechanisms. Though this remains to be formally tested, one scenario in which El Tor *Vibrio cholerae* defends against phage could be a reliance of the non-lethal depletion of nucleotides by the *avcID* system (8, 9) at LCD and the CBASS system at HCD, both of which are encoded in VSP-1. Although the molecular mechanism of CBASS are now well characterized (55), the regulation of such systems and their contribution to bacterial evolution and environmental adaptation is just beginning to be described. CBASS are widely conserved in bacteria (19) and whether QS regulation of such systems is commonplace should be further investigated.

## Materials and Methods

### Growth conditions and media

All strains of *V. cholerae* and *E. coli* were grown in LB broth-Miller media (NEOGEN) at 35° C with aeration, unless otherwise stated. When noted, antibiotic selection was utilized at the following concentrations: streptomycin (50 µg/mL), chloramphenicol (10 µg/mL when used alone or 5 µg/mL when used with another antibiotic), kanamycin (100 µg/mL when used alone; 50 µg/mL when used with another antibiotic) and tetracycline (5 µg/mL). IPTG was used at 6.25-10 μM as indicated. *E. coli* BW29427, a diaminopimelic acid (DAP) auxotroph, was additionally supplemented with 300 µg/mL DAP and used for the conjugative transfer of vectors and cosmids to all *V. cholerae* strains presented in this work.

### Cloning and strain construction

All gene deletions from the *V. cholerae* genome were performed using the vector pKAS32 (56). Deletion constructs were cloned using a three-piece Gibson Assembly composed of ∼1 kb homologous sequences both 5’ and 3’ of the genomic region to be removed and linear pKAS32 double digested with KpnI and SacI. Mutants were obtained through allelic exchange (56) and verified by Sanger sequencing. All mutants utilized in this study were complete deletions of the genomic regions of interest except for Δ*vspR* where the *vspR* codons 1, 2, and 6 where all mutated into stop codons. The pP_CBASS_::*lux* plasmid was generated by amplification of 913 n.t. upstream of *capV* and Gibson assembly into BamHI/SpeI digested pBBR-lux (57) using the primers Lux_CBASSpr_FW and Lux_CBASSpr_RV, 5’.

### Metabolic Growth Assays

Using an inoculating loop, overnight cultures were applied as a single streak on the surface of agar plates and incubated at 35° C for 24 h. Images were taken of plates using an iPhone. For the casein hydrolysis protease assay, milk agar plates were prepared according to (4) and contained 20.0 g/L dry skim milk and 9.2 g/L brain-heart infusion and 15 g/L agar. Blood agar plates were prepared according to (58) using Mueller Hinton Broth in 1.5% agar with 5% sheep blood. Citrate minimal medium agar was prepared according to (4). For comparing growth on MacConkey agar, an overnight culture of each strain was normalized to an OD_600_ of 0.5 and serial diluted ten-fold in PBS to 10^−7^. 2.5 µl of each dilution was plated on both LB and MacConkey agar plates and incubated ∼16 h at 35° C and imaged with an iPhone. For the Voges-Proskauer (VP) Assay, each strain was inoculated in 3 mL of Methyl Red - Voges-Proskauer (MR-VP) broth medium (4) from a plate and incubated overnight. Then, 130 µL of 5% (w/v) α-naphthol and 43 μl of 1M potassium hydroxide was added to 1mL aliquots of the overnight cultures and allowed color to develop over 48 h at room temperature.

### Infant Mouse Competition Assay

Infant mice were infected as described previously (59, 60). Briefly, three-to five-day old CD-1 neonate mice (Charles River, Wilmington, MA) were orogastrically inoculated with approximately 10^6^ total CFU 2 h after separation from dam mice. ΔVSP-1/2 was co-inoculated 1:1 with a Δ*lacZ* C6706str2 strain to allow for differentiation by blue-white screening upon recovery. Mice were maintained at 30°C until euthanasia 20 h post inoculation. To enumerate *V. cholerae* CFU, intestinal segments (small intestine and the large intestine plus cecum) were homogenized, serially diluted, and plated on LB + 0.1 mg/mL streptomycin, 0.08 mg/mL 5-bromo-4-chloro-3-indolyl-β-D-galactopyranoside (X-Gal). Blue and white colony counts were used to determine the competitive index (C.I.) for ΔVSP-1/2 using the following equation: C.I. = (CFU ΔVSP-1/2_intestine_ / CFU Δ*lacZ*_intestine_)/(CFU ΔVSP-1/2_inoculum_ / CFU Δ*lacZ*_inoculum_). The fitness of the Δ*lacZ* strain was previously shown to have no colonization defect relative to the parental C6706str2 (59). All animal experiments in this study were approved by the Institutional Animal Care and Use Committee at Michigan State University.

### Polymyxin B IC_50_

For the polymyxin B resistance assay, 2 mL LB cultures were incubated for 16 h at 35° C with aeration. Cultures were then diluted 1:100 in fresh LB, aliquoted into 96-well plates (COSTAR), and challenged with a four-fold serial dilution of polymyxin B (Sigma) from 55.5 μg/mL to 0.014 μg/mL. Plates were incubated for 20 h without aeration at 35° C and the culture absorbance was measured at 600 nm using an Envision 2105 Multimode Plate Reader (Perkin-Elmer) and % growth for each biological replicate was calculated by dividing the absorbance 600 nm of polymyxin B treated wells by the absorbance of the untreated control well. The reported mean % growth and standard deviation were calculated from three biological replicates for each polymyxin B concentration. IC_50_ were calculated using a non-linear regression analysis performed using GraphPad Prism version 9.5.0.

### Sulfamethoxazole IC_50_ and Challenges

Overnight cultures were grown in the presence of streptomycin unless pLAFR derived cosmids were being maintained, in which case tetracycline was supplemented in place of streptomycin. Sulfamethoxazole (10 mg/mL) stocks were prepared in DMSO, diluted in LB to concentrations 15 x greater than the final desired concentration, and multichannel pipetted into clear polystyrene 96-well plates (COSTAR) in 10 µL aliquots. Overnight cultures were diluted 1:10,000 in fresh media supplemented with the same antibiotic selection as the overnight media. One hundred and forty µL of the diluted cultures were multichannel pipetted into 96-well plates preloaded with trimethoprim or sulfamethoxazole. Plates were wrapped in parafilm and incubated at 35° C with aeration for 24 h. Culture absorbance was measured at 600 nm using an Envision 2105 Multimode Plate Reader (Perkin-Elmer). % growth for each biological replicate was calculated by dividing the mean absorbance of technical replicate treatment wells (N = 2 to 4) by the mean absorbance of technical replicate untreated control wells (N = 2 to 4). The reported mean % growth and standard error mean were calculated from three biological replicates for each concentration. IC_50_ were calculated using a non-linear regression analysis performed using GraphPad Prism version 9.5.0.

### Growth curves and cGAMP quantification by UPLC-MS/MS following Sulfamethoxazole challenge

Overnight cultures were started in triplicate from freezer stocks and grown in LB supplemented with streptomycin. Cultures were inoculated 1:1,000 in 50 mL fresh media containing selection in 125 mL flasks and grown for 15 minutes before being divided equally into two sister cultures. Sister cultures were grown for one additional hour before being sampled to assess culture growth by measuring absorbances at 600 nm and CFU enumeration by serial dilution plating. Immediately following the initial sampling, sister cultures were challenged with either 100 µg/mL SMX or a DMSO vehicle control and further sampling continued at ∼30 minute intervals for the duration of the experiment. We elected to challenge these strains with 100 µg/mL SMX as this was sufficient to induce CBASS dependent SMX sensitivity in C6706 and below the IC_50_ of ΔVSP-1 (Figs. 2A, S2A & Table S1), which has an analogous SMX resistance phenotype to the Δ*capV* strain (Figs. 2D & S2F).

For the purposes of measuring intracellular cGAMP additional aliquots of Δ*capV* sister cultures (+/−100 µg/mL sulfamethoxazole) were removed at all time points during this experiment and similarly analyzed as previously described (10). One-milliliter culture aliquots were pelleted at 15k x g in microcentrifuge tubes for 1 minute, supernatants were removed by aspiration, and pelleted cells were suspended in 200 µL of ice-cold extraction buffer (acetonitrile, methanol, HPLC-grade water, formic acid (2:2:1:0.02, v/v/v/v)), and stored at −20° C overnight. Extracts were centrifugation at 15k x g for 1 minute, to remove cellular debris, and the resulting clarified extracts were transferred to new a microcentrifuge tube and dried in a SpeedVac. Desiccated extracts were dissolved in 100 µL of HPLC-grade water and loaded into glass sample vials for UPLC-MS/MS analysis using an Acquity Ultra Performance LC system (Waters) coupled with a Quattro Premier XE mass spectrometer (Waters). Chromatography and multiple reaction monitoring parameters performed as previously described (10). A cGAMP standard curve was generated using a two-fold serial dilution of cGAMP (Axxora) in HPLC-grade water spanning 1.9-125 nM. Intracellular concentrations of cGAMP were calculated by dividing the total moles of cGAMP in a sample by the product of the enumerated CFU in each sample and a standard cell volume of 6.46 × 10^−16^ L (10, 61).

### Growth curves of sulfamethoxazole treated quorum sensing and CBASS mutants

Overnight cultures were diluted to an OD_600_ of 0.01 in 6 mL fresh LB medium, recovered for 1h at 37° C with aeration, and split into paired test tubes. Within a pair of cultures, one was challenged with 100 µg/mL sulfamethoxazole, dissolved in DMSO, while the second culture was challenged with DMSO vehicle control. Cultures were incubated at 37° C with aeration and the culture OD_600_ was measured at the times presented for the duration of each experiment. The mean and standard deviation of biological triplicate samples for all strains and treatments is reported.

### Time course gene expression using RT-qPCR

Biological duplicate overnight cultures were started from freezer stock in LB and back diluted 1:10,000 into 250 mL LB in 1 L flasks and grown at 35° C with aeration. Cultures were sampled 2, 2.5, 3, 4, and 5 h post inoculation in 50, 30, 1, 0.5, and 0.5 mL aliquots, respectively, and cells were pelleted by centrifugation. Cell pellets were suspended in 1 mL TRIzol^TM^ Reagant (Thermo Fischer) and RNA was purified following manufacturer’s specifications. Following manufacturer recommendations, 5 µg of RNA was treated with TURBO^TM^ Dnase (Ambion) and cDNA was generated using SuperScript^TM^ III (Thermo Fischer). SYBR^TM^ Green PCR Master Mix (Thermo Fischer) was used according to the manufacture’s recommendations in 25 µL reactions containing 6.25 ng cDNA template (no reverse-transcription controls used 6.25 ng DNAse-treated RNA template) and a final primer concentration of 100 nM. qRT-PCRs reactions were performed in technical duplicate using a StepOnePlus real-time PCR system (Thermo Fisher Scientific). Gene expression was calculated using ΔCT relative to the *gyrA* housekeeping gene and comparative ΔΔCT was determined by comparison of each time point to the ΔCT of the initial 2 h sample (∼0.05 OD_600_).

### Gene expression in response to exogenous autoinducers by RT-qPCR

Colonies of Δ*csqA*Δ*luxS V. cholerae* were inoculated into LB and incubated with aeration at 30° C for 16 hours. Each culture was then diluted 1:500 – once into fresh LB supplemented autoinducers (5 µM CAI-1 and 5 µM AI-2) and once into fresh LB without autoinducers. These cultures were incubated with aeration at 30° C for 2 hours, at which point the OD_600_ measurements for all cultures were confirmed to be similar (OD_600_ = 0.4-0.45). Cells were lysed in TRIzol^TM^ Reagent (Invitrogen) then RNA was extracted using the Direct-zol RNA Microprep kit (Zymo Research). RNA was then treated with 0.5 µl TURBO^TM^ DNase (Invitrogen) at 37° C for 90 minutes, then an additional 0.5 µl TURBO^TM^ DNAse was added and the samples were incubated for an additional 90 minutes at 37° C. cDNA was synthesized from total RNA using the SuperScript^TM^ III Reverse Transcriptase kit (Invitrogen). For each sample, RT-qPCR was performed on 50 ng of cDNA using the SYBR^TM^ Select Master Mix kit (Applied Biosystems) and 250 nM of each primer. Expression levels were calculated for each target gene by normalizing to the housekeeping gene *recA*, then the relative fold expression was calculated by comparing the target gene expression in the presence of autoinducers to the absence of autoinducers and reported as the mean of measurements obtained from three biological replicates.

### Luminescent reporter assays

To assess QS induction of CBASS transcription, overnight cultures of Δ*vpsL*, Δ*vpsL*Δ*hapR*, and Δ*vpsL*Δ*luxO* (35) containing pP_CBASS_::*lux* (pKAD1) inoculated from individual colonies (n=3) grown in LB in glass test tubes were back diluted 1:1,000 in 3 mLs of LB with chloramphenicol and grown with shaking at 35° C. At two hours, to measure low cell density, and 20 hours, to measure high cell density, the bioluminescence and OD_600_ of 200 mL of each culture was transferred to a solid black 96-well plate and quantified on an Envision 2105 Multimode Plate Reader (Perkin-Elmer). Relative light units were determined by dividing the bioluminescence by the OD_600_, and each was normalized to the mean relative light units of the locked high cell density Δ*vpsL*Δ*luxO* mutant at the corresponding time point.

### pHapR and pVSP-1 in E coli

Three independent 2 mL overnight cultures of *E. coli* DH10B with pP_CBASS_::*lux* (pKAD1) with pVector2 (pEVS141) (62) or pHapR (pSLS13) (63) grown in LB with chloramphenicol (5 μg/mL) and kanamycin (50 μg/mL) were back diluted 1:100 in the same media. 150 μL of each culture was placed in a solid black 96-well plate and incubated without shaking at 35° C. After 1.5 hours, HapR production was induced with 6.25 μg/mL IPTG, and bioluminescence and OD600 were measured on an Envision 2105 Multimode Plate Reader (Perkin-Elmer) hourly.

### Phage Challenge Assay

Overnight cultures of *E. coli* DH10B with pVector (pLAFR) (64) or pVSP-1 (10) combined with pVector2 (pEVS141) (62) or pHapR (pSLS13) (63) were grown in 2 mL LB with 50 μg/mL kanamycin and 5 μg/mL tetracycline overnight at 35° C with shaking. Each culture was back diluted 1:1,000 in 2 mL of LB with the same antibiotics + 10 µM IPTG and grown 2-3 hours with shaking at 35° C before addition of T2 phage at the indicated MOI followed by overnight growth with shaking at 35° C. The OD_600_ was measured the following day (∼20 hours) on an Envision 2105 Multimode Plate Reader (Perkin-Elmer), and the OD_600_ of each culture was normalized to the uninfected control and reported as percent growth.

## Figure Legends

**Figure S1:**
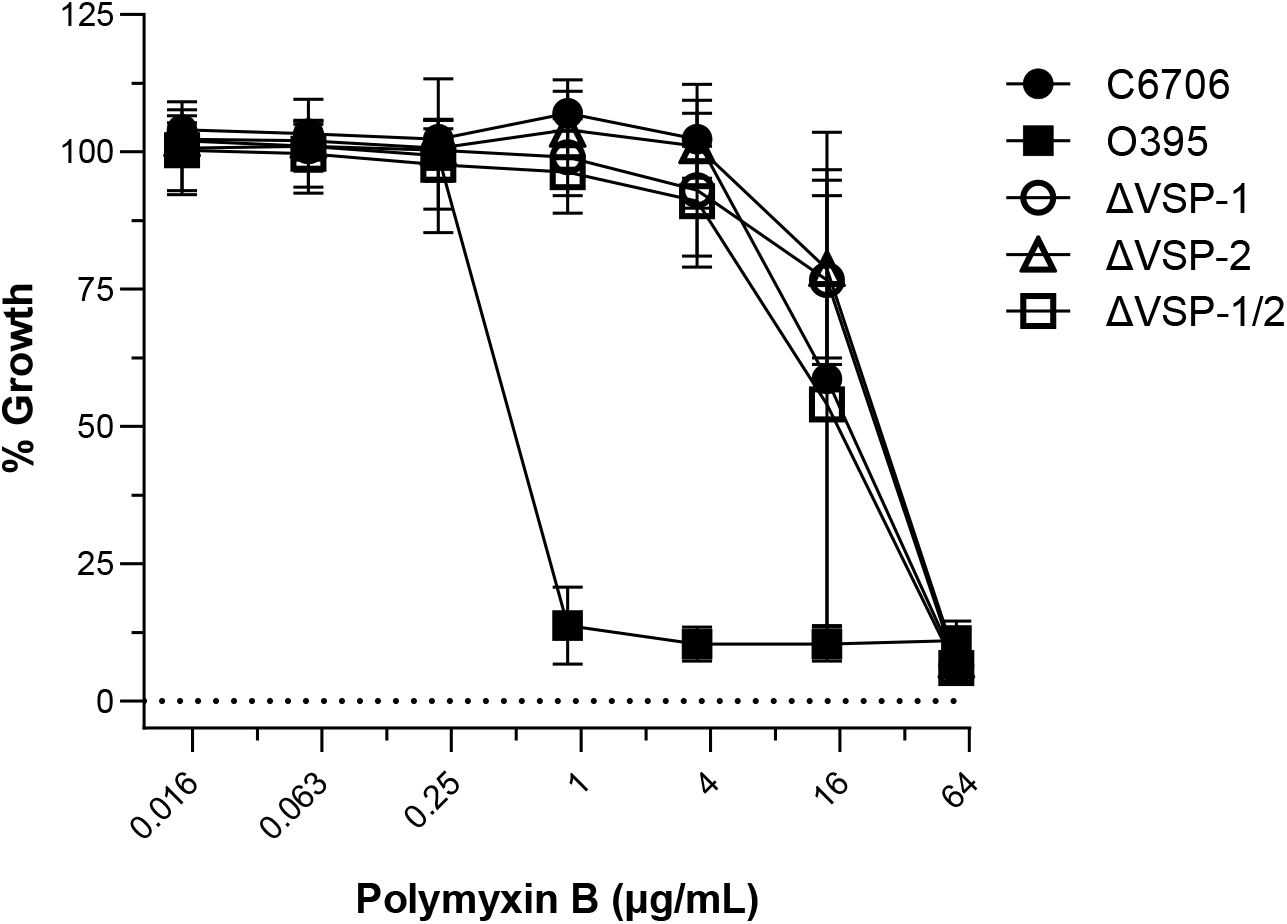
VSP-1 & -2 do not contribute to C6706 resistance to the antimicrobial peptide polymyxin B. 20-hour planktonic antibiotic sensitivity assay performed using a polymyxin B concentration gradient. % Growth reported as (OD_600_ polymyxin B treated / OD_600_ untreated) after 20 hours. Dotted line indicates 0% Growth. N = 3 biological replicates for all data points. Error bars represent standard deviation. IC_50_ for all strains are presented in Supplementary Table 1.

**Figure S2:**
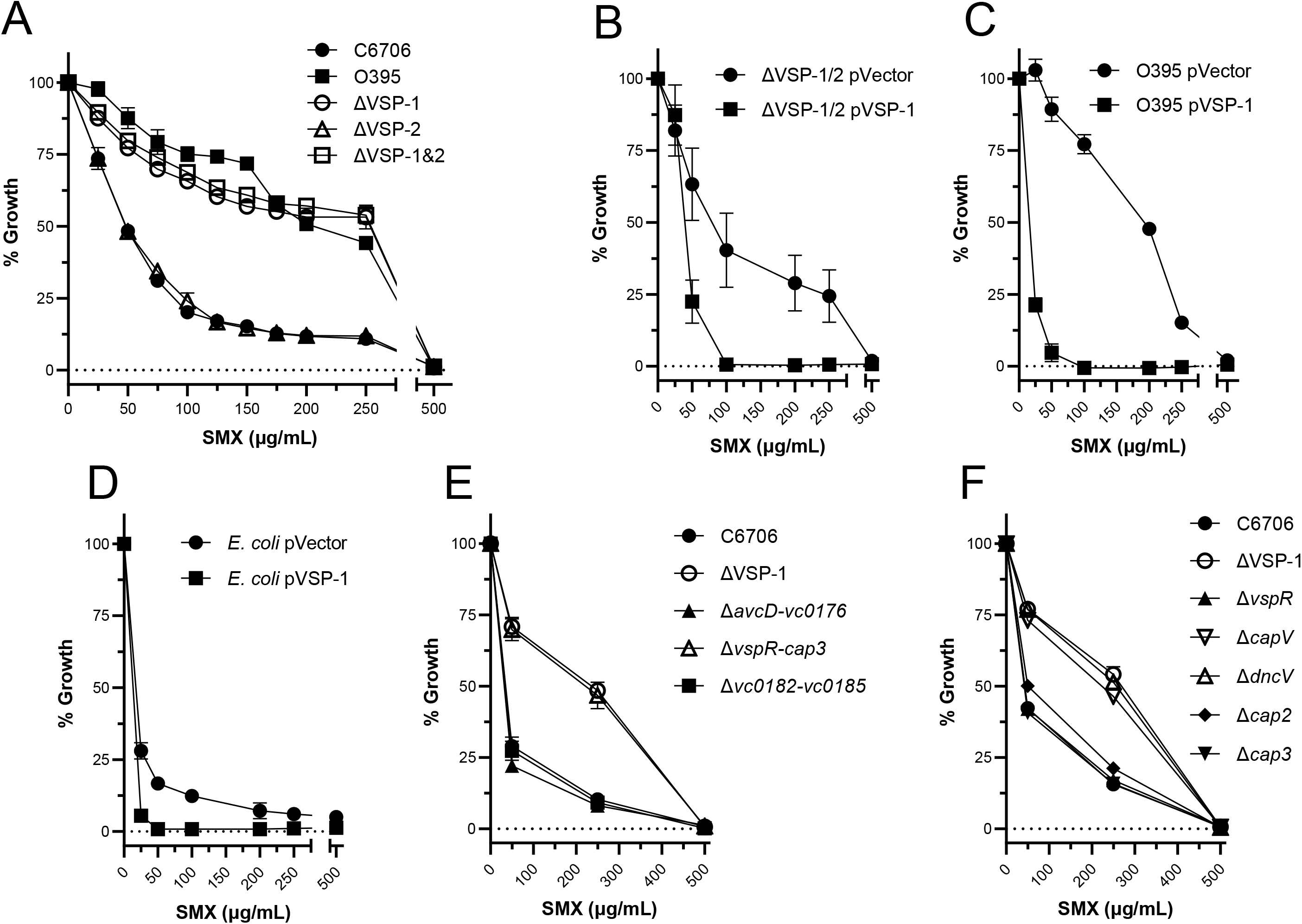
Scatter plots showing that VSP-1 encoded CBASS is responsible for *V. cholerae* biotype specific SMX sensitivity. (**A-F**) 24-hour planktonic antibiotic sensitivity assays performed in a variety of SMX concentration gradients. Scatter plots represent are the same data presented in heatmap form in (Figs. 2A, 2B, & 2D). N = 3 biological replicates and error bars represent standard error of the mean. IC_50_ for all strains in (**A**) are presented in Supplementary Table 1.

**Figure S3:**
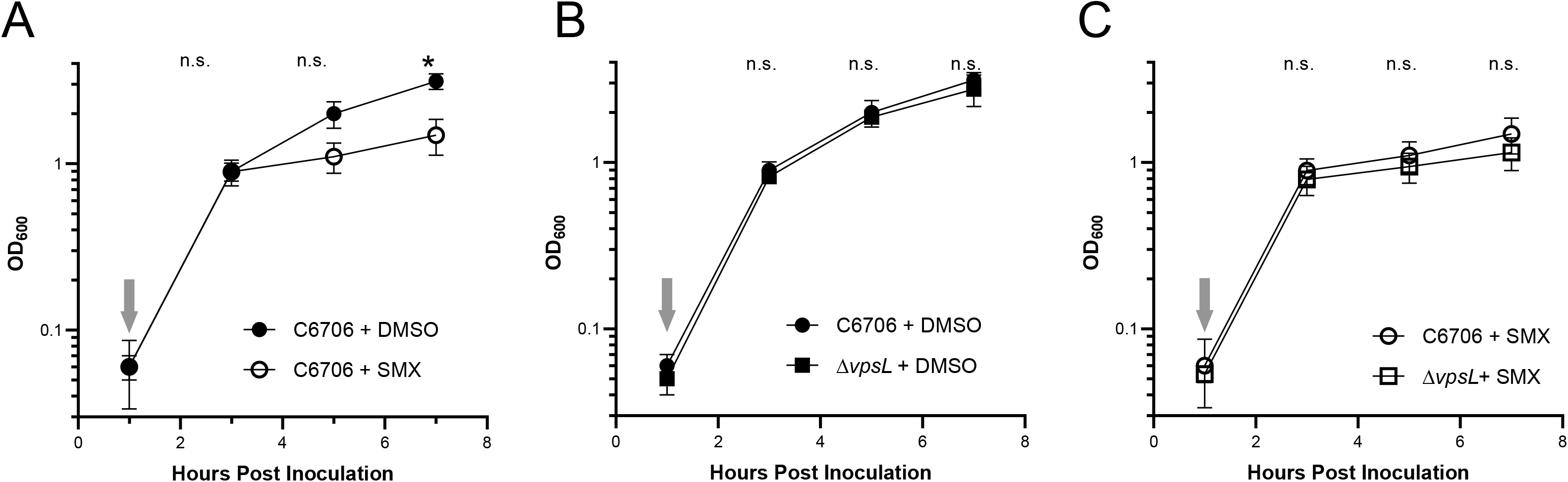
Lack of biofilm formation does not alter *V. cholerae* C6706 sensitivity to SMX. Growth curves of (**A**) *V. cholerae* C6706 treated without (+ DMSO) and with 100 µg/mL SMX (+ SMX), and C6706 and Δ*vpsL* (**B**) untreated (+DMSO) and (**D**) treated (+SMX) with 100 µg/mL SMX. Grey arrows indicate addition of 100 µg/mL SMX or DMSO, approximately 1-hour after cultures were inoculated. N = 3 biological replicates and error bars represent standard deviation. For the purposes of statistical analysis, Δ*vpsL* data presented in (**B**) and (**C**) are also presented in (Fig. 3B). Statistical significance calculated using an unpaired *t* test with the Holm-Šídák method (*P < 0.05), n.s. = not significant.

**Figure S4.**
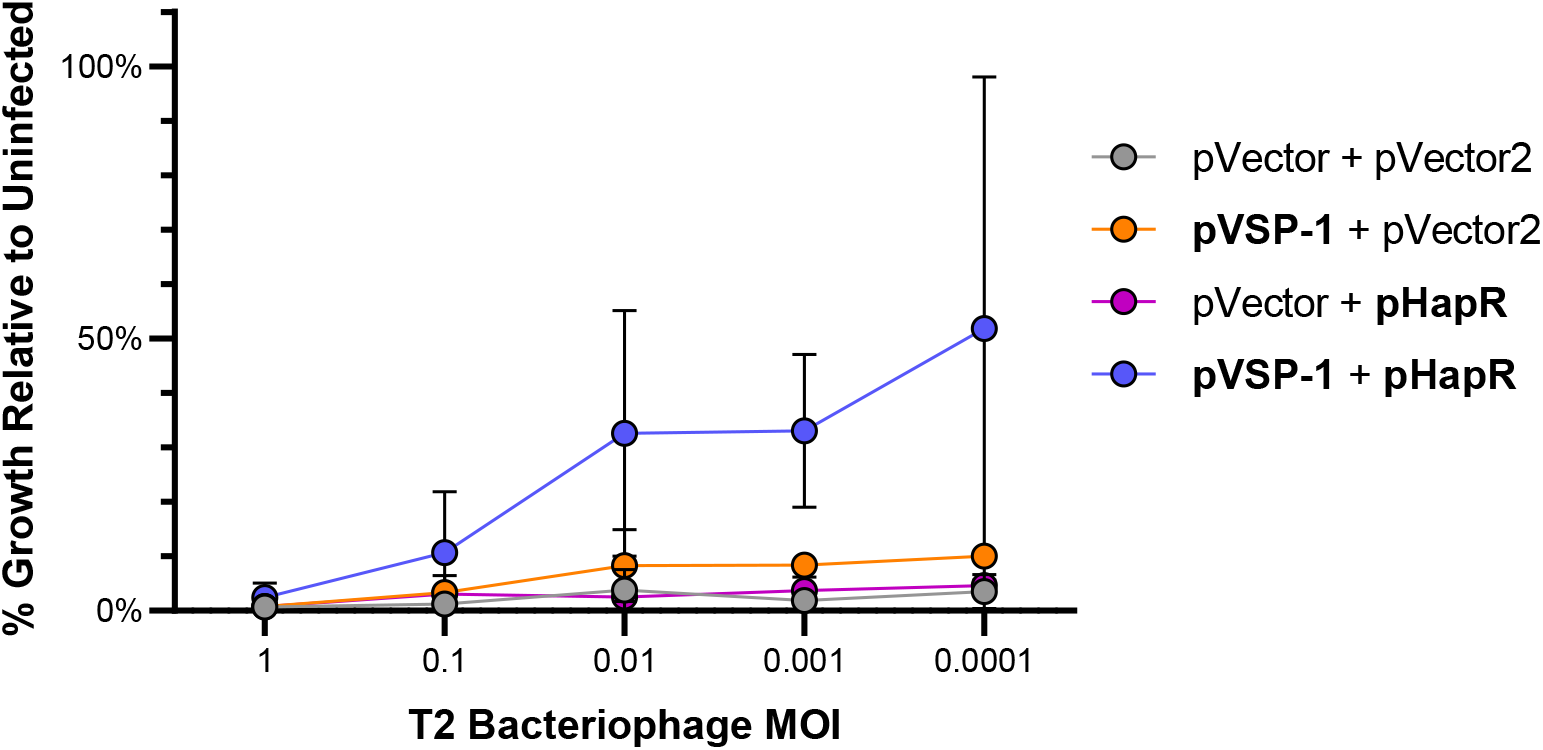
T2 phage infection graphs. Growth of *E. coli* containing either pHapR induced with 10 μM IPTG and pVSP-1 with their associated vector controls after overnight growth with T2 phage. Data represent mean percent growth (OD_600_) at the specified MOI calculated relative to control uninfected cultures. N = 3 biological replicates and error bars represent standard deviation. Data presented in heat map form in (Fig. 5A).

**Supplementary Table 1:** *V. cholerae* Antibiotic Sensitivity.

| Strain | Antibiotic | IC<sub>50</sub> (µg/mL) | R<sup>2</sup> |
| --- | --- | --- | --- |
| C6706 | Polymyxin B | 18.4 | 0.77 |
| O395 | Polymyxin B | 0.6 | 0.85 |
| ΔVSP-1 | Polymyxin B | 22.1 | 0.84 |
| ΔVSP-2 | Polymyxin B | 24.8 | 0.81 |
| ΔVSP-1/2 | Polymyxin B | 14.5 | 0.84 |
| C6706 | Sulfamethoxazole | 36.5 | 0.95 |
| O395 | Sulfamethoxazole | 230.7 | 0.82 |
| ΔVSP-1 | Sulfamethoxazole | 184.7 | 0.85 |
| ΔVSP-2 | Sulfamethoxazole | 38.0 | 0.95 |
| ΔVSP-1/2 | Sulfamethoxazole | 207.2 | 0.85 |

**Supplementary Table 2.**
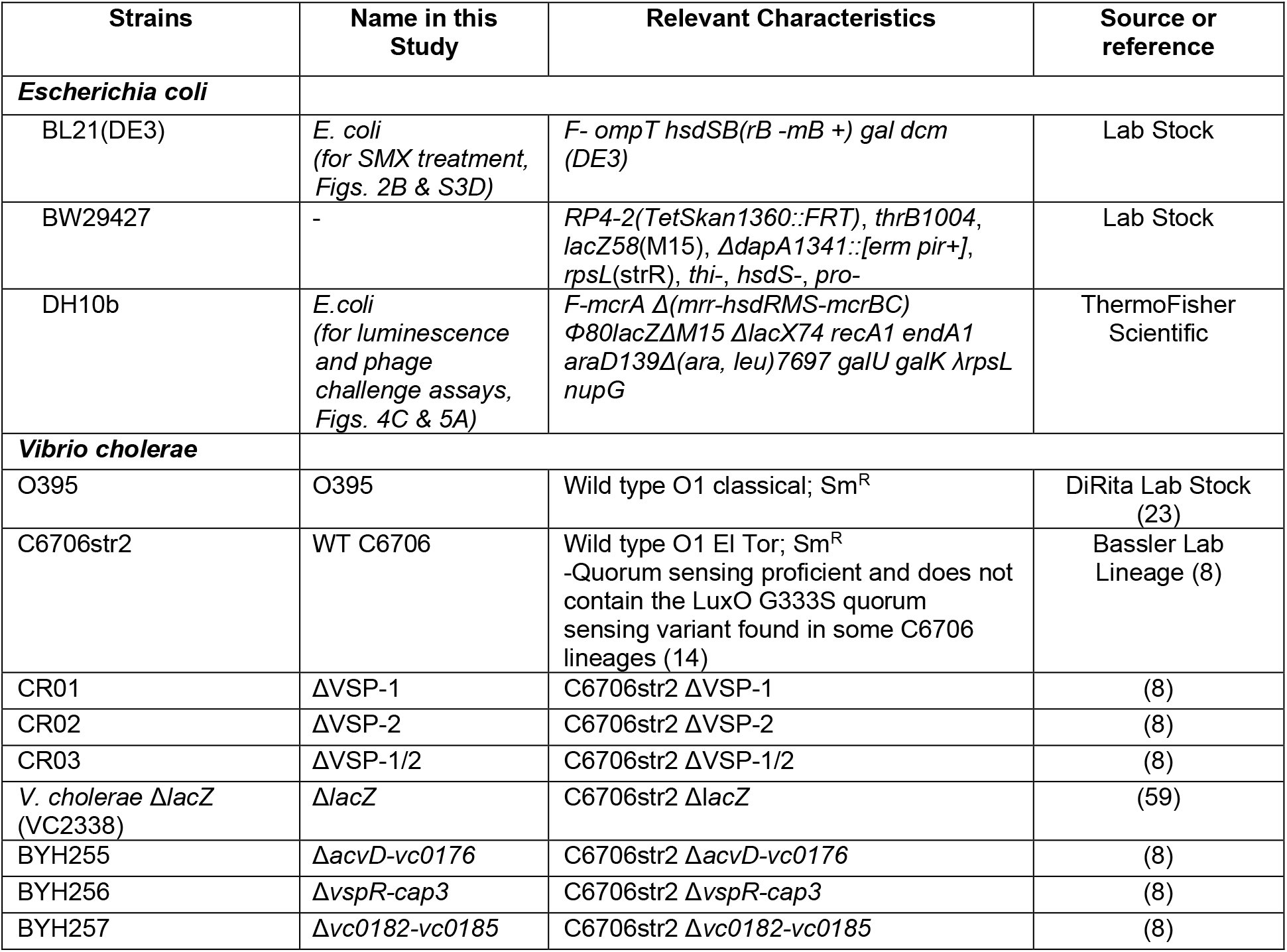
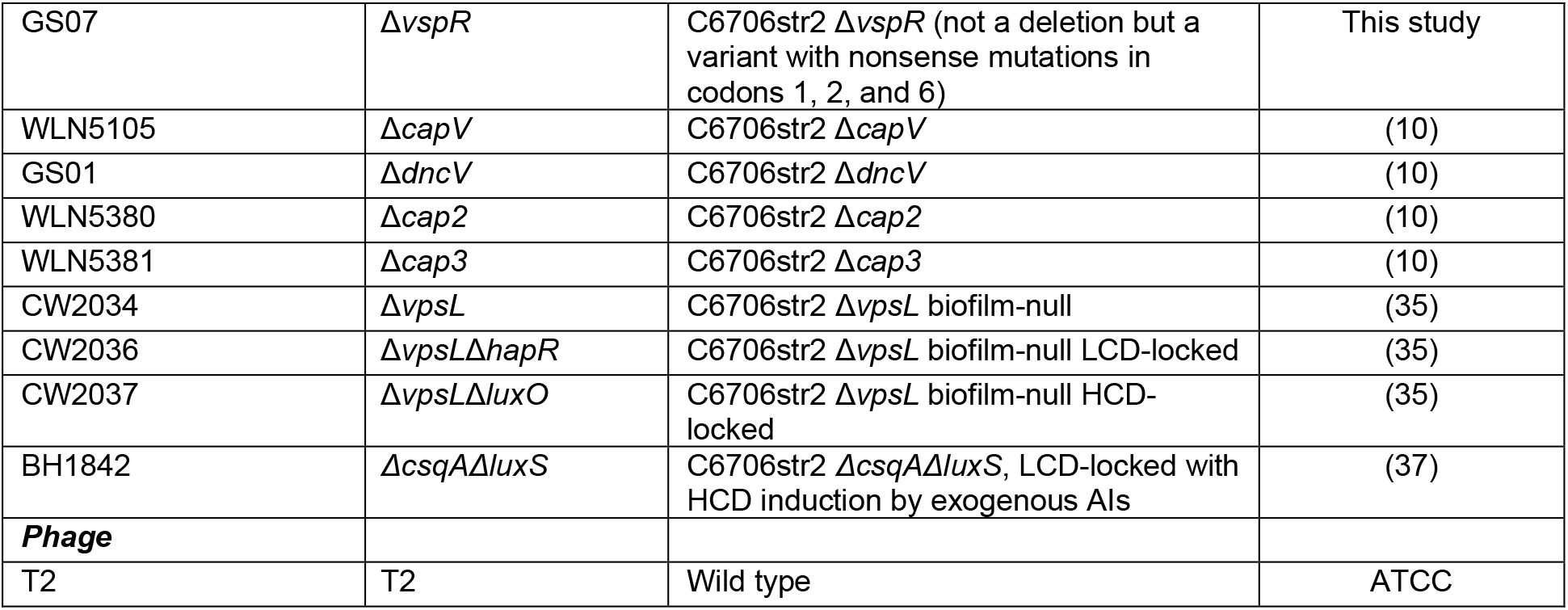
Bacterial strains used in this study.

**Supplementary Table 3:** Plasmids used in this study.

| Plasmids | Name in this Study | Relevant characteristics | Source or Reference |
| --- | --- | --- | --- |
| pLAFR | pVector | pLAFR cosmid; Tet<sup>r</sup> | (64) |
| pCCD13 | pVSP-1 | pLAFR containing VSP-1; Tet<sup>r</sup> | (10) |
| pBBR-lux | - |  | (57) |
| pKAD1 | pP<sub>CBASS</sub>::lux | pBBRlux containing 913bp 5' of *capV* locus (*i.e.*, CBASS promoter) to regulate luciferase expression | This study |
| pGBS54 | - | pKAS32; Amp<sup>r</sup> containing homologous region for introducing *vspR* nonsense mutations (creating strain GS07) | This study |
| pSLS13 | pHapR | *hapR* allele cloned into pEVS143 under P<sub>tac</sub> | (63) |
| pEVS141 | pVector2 | Promoterless pEVS143 vector control | (62) |

**Supplementary Table 4:**
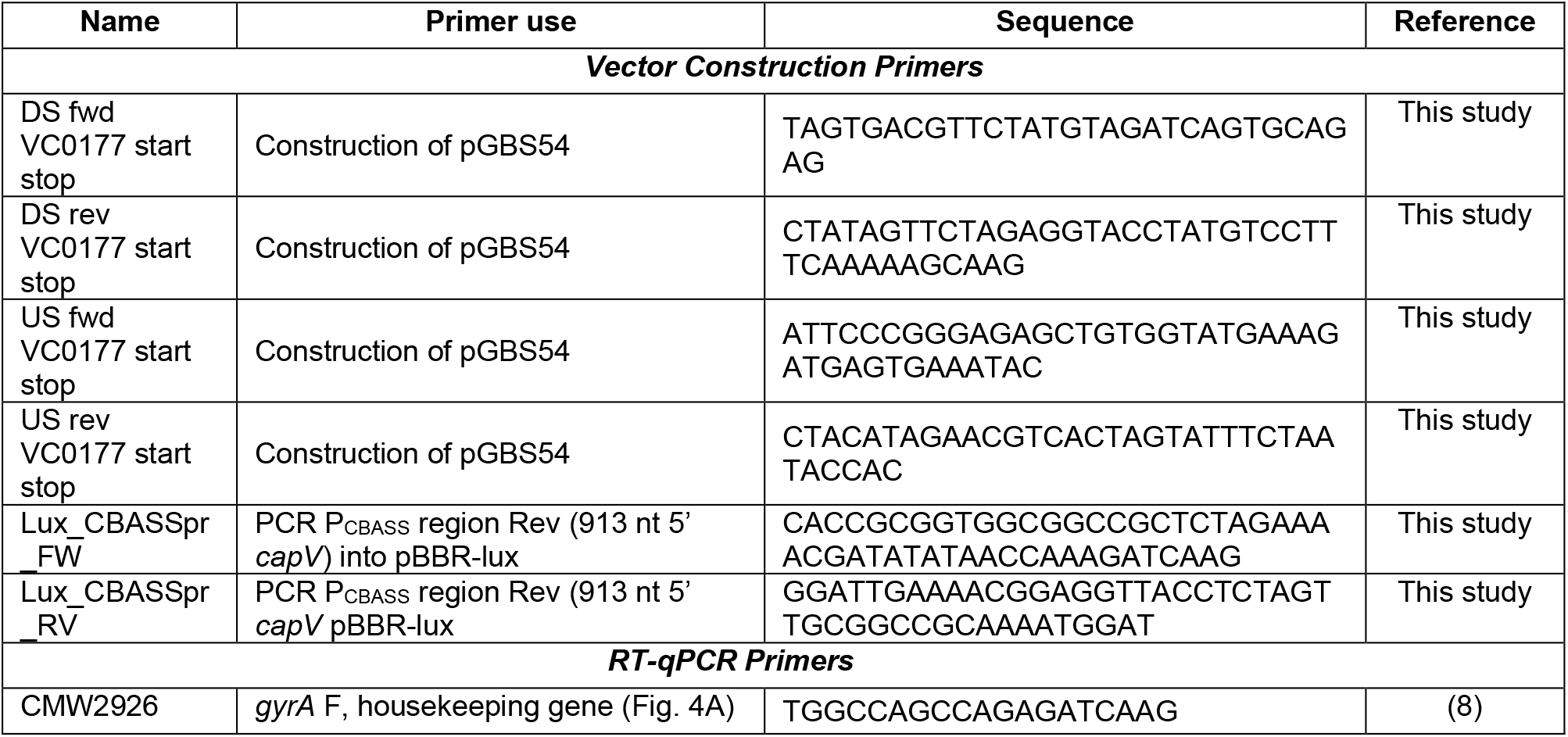
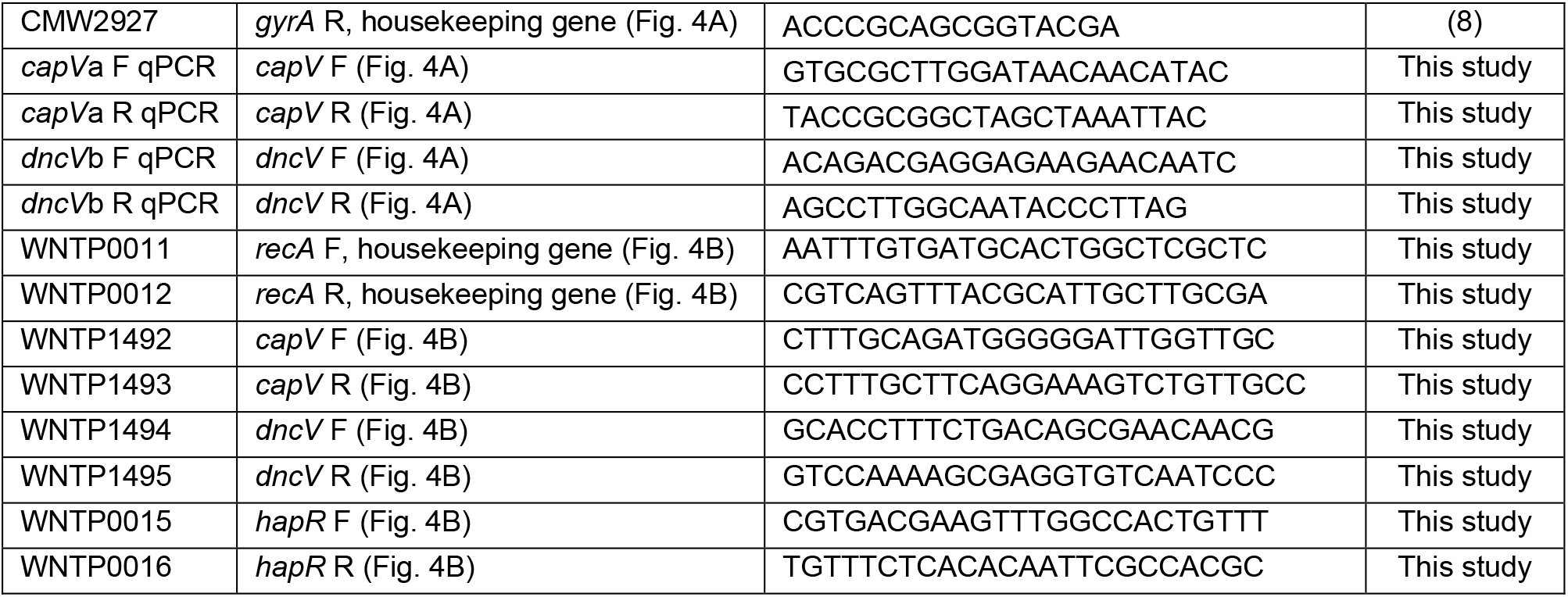
Oligonucleotides used in this study.

